# Serotonergic drugs inhibit CHIKV infection at different stages of the cell entry pathway

**DOI:** 10.1101/2020.03.24.005066

**Authors:** Ellen M. Bouma, Denise P.I. van de Pol, Ilson D. Sanders, Izabela A. Rodenhuis-Zybert, Jolanda M. Smit

**Affiliations:** Department of Medical Microbiology and Infection Prevention, University Medical Center Groningen, University of Groningen, Groningen, The Netherlands

## Abstract

Chikungunya virus (CHIKV) is an important re-emerging human pathogen transmitted by mosquitoes. The virus causes an acute febrile illness, chikungunya fever, which is characterized by headache, rash and debilitating (poly)arthralgia that can reside for months to years after infection. Currently, effective antiviral therapies and vaccines are lacking. Due to the high morbidity and economic burden in the countries affected by CHIKV, there is a strong need for new strategies to inhibit CHIKV replication. The serotonergic drug, 5-nonyloxytryptamine (5-NT), was previously identified as a potential host-directed inhibitor for CHIKV infection. In this study, we determined the mechanism of action by which the serotonin receptor agonist 5-NT controls CHIKV infection. Using time-of-addition and entry bypass assays we found that 5-NT predominantly inhibits CHIKV in the early phases of the replication cycle; at a step prior to RNA translation and genome replication. Intriguingly, however, no effect was seen during virus-cell binding, internalization, membrane fusion and gRNA release into the cell cytosol. Additionally, we show that the serotonin receptor antagonist MM also has antiviral properties towards CHIKV and specifically interferes with the cell entry process and/or membrane fusion. Taken together, pharmacological targeting of 5-HT receptors may represent a potent way to limit viral spread and disease severity.

**Importance:** The rapid spread of mosquito-borne viral diseases in humans puts a huge economic burden on developing countries. For many of these infections, including Chikungunya virus (CHIKV), there are no specific treatment possibilities to alleviate disease symptoms. Understanding the virus:host interactions that are involved in the viral replication cycle is imperative for the rational design of therapeutic strategies. In this study, we discovered an antiviral compound and elucidated the mechanism of action and propose serotonergic drugs as potential host-directed antivirals for CHIKV.

## Introduction

Chikungunya fever is an important re-emerging mosquito-borne human disease caused by Chikungunya virus (CHIKV). Over the past decade, the virus has continued to spread throughout the Americas and Asia thereby infecting millions of people (1). Chikungunya fever is characterized by fever, headache, rash, and myalgia. A potential long-lasting and debilitating feature of CHIKV infection is the onset of (poly)arthralgia and/or polyarthritis which can last months to years after infection (2, 3). Roughly 85% of all infected individuals develop chikungunya fever of which approximately 30-40% develop long lasting (poly)arthralgia/arthritis (1, 4). Consequently, CHIKV has a high morbidity and economic burden in the countries affected especially since there are no vaccines nor antiviral therapies available.

Antiviral therapies against CHIKV should be designed with the aim to lower viral burden and/or to prevent the onset of chronic disease. There are two classes of antivirals: 1) direct-acting drugs, which target the virus itself and 2) host-directed drugs, which target cellular factors important for the replication cycle of the virus (5–7). An advantage of direct-acting antivirals is that these are more specific, however, viral resistance is often quickly obtained (8). Host-directed antivirals, on the other hand, are less specific and may cause more side-effects yet the development of viral resistance is greatly reduced (9). To identify novel host-directed antivirals, it is imperative to understand the dynamic and temporal interactions of the virus with the host during infection.

To initiate infection, CHIKV interacts with cellular receptors expressed at the plasma membrane. Among others, the cell adhesion molecule Mxra8 and N-sulfated heparan sulfate have been proposed as putative receptors for CHIKV, thereby facilitating virus internalization via clathrin-mediated endocytosis (6, 10). The acidic lumen of the early endosome subsequently triggers conformational changes in the E2 and E1 viral spike glycoproteins leading to E1-mediated membrane fusion (11, 12). Thereafter, the viral nucleocapsid is dissociated by an as yet ill-understood process and the positive-sense RNA is translated to form the non-structural proteins of the virus. These non-structural proteins interact with multiple cellular factors to facilitate 1) RNA replication, 2) translation of structural proteins from the viral subgenomic mRNA, and 3) production of new genomic RNA. The structural proteins E1 and a precursor E2 are translocated to the ER where heterodimerization of E1/E2 occurs. Maturation of the E1/E2 viral spike complex occurs via transit through the cellular secretory pathway. Progeny genomic RNA interacts with newly produced viral capsid proteins to form a nucleocapsid which is transported to the plasma membrane where virion assembly and budding occurs (13, 14).

Serotonin (5-hydroxytryptamine; 5-HT) receptors are expressed at the plasma membrane and known to facilitate or alter the infectivity of different classes of viruses (15– 17). Most of the 5-HT receptors are G-protein coupled receptors and regulate important physiological functions and signaling pathways, including the cycling adenosine monophosphate (cAMP), calcium and phosphatidylinositol pathways (18). There are multiple subtypes of 5-HT receptors and these are divided into 7 families (19). The 5-HT_2_ receptor family has been described to facilitate cell entry of JC polyomavirus (20) and 5-HT_1_ is suggested to be involved in HIV-1 replication (21). Furthermore, the infectivity of multiple RNA viruses were found to be controlled by 5-HT receptor agonists and antagonists (9, 22–24). Indeed, reovirus and CHIKV infectivity was found reduced in the presence of the 5-HT receptor agonist 5-nonyloxytryptamine (5-NT), which is described as a specific 5-HT_1B_/5-HT_1D_ receptor agonist though it has also low level affinity to other 5-HT receptor families (22, 25). 5-NT was shown to interfere with reovirus intracellular transport and disassembly kinetics during cell entry (22). Opposed to the agonist, the 5-HT receptor antagonist methiothepin mesylate (MM) increased reovirus infectivity. However, it is yet unclear whether the mechanism of action of 5-HT receptor stimulation with this 5-HT receptor agonist and antagonist is the same for CHIKV.

In this study, we confirmed the antiviral properties of 5-NT and unraveled the mode of action in CHIKV infection. Also, and in contrast to that observed for reovirus, we found an antiviral effect of the serotonin receptor antagonist MM towards CHIKV. We show that the serotonergic drugs 5-NT and MM target distinct steps during CHIKV cell entry and conclude that targeting 5-HT receptors may be a novel strategy to alleviate CHIKV disease.

## Results

### 5-NT strongly inhibits CHIKV infection and virus particle production in U-2 OS cells

The effect of 5-NT on CHIKV infectivity was analyzed in human bone osteosarcoma epithelial U-2 OS cells as epithelial cells are natural targets during human CHIKV infection (29, 30). Also, these cells were used by Mainou and co-workers who previously identified 5-NT as an inhibitor of CHIKV and reovirus infection (22). First, the mRNA expression levels of 10 distinct 5-HT receptor subtypes were determined in U-2 OS cells. We confirmed expression of 8 distinct 5-HT receptor subtypes including the 5-HT_1B_ and 5-HT_1D_ receptor to which 5-NT binds with high affinity (Fig 1A). Next, we assessed the cellular cytotoxicity of 5-NT in U-2 OS cells by MTT assay and revealed no significant cytotoxicity up to a concentration of 5µM 5-NT (Fig 1B). The highest dose in our infectivity experiments was therefore set at 5µM 5-NT. 5-NT was dissolved in DMSO and the final DMSO concentration was below 1% in all experiments. Then, we analyzed CHIKV-LR infectivity and infectious virus particle production in U-2 OS cells in the presence of increasing concentrations of 5-NT. Cells were pretreated with increasing doses of 5-NT or NH_4_Cl, a lysosomotropic agent known to neutralize the endosomal pH and thereby inhibiting the membrane fusion activity of CHIKV, for 1 h and infected with CHIKV-LR 5’GFP at MOI 5. At 20 hpi, cells and supernatants were collected and analyzed for GFP expression by flow cytometry and the production of infectious virus particles by plaque assay, respectively. A clear dose-dependent reduction in the number of CHIKV-infected cells was observed (Fig 1C). Importantly, the vehicle control DMSO had no significant effect on the number of infected cells (Fig 1C). CHIKV infection was reduced from 50.8% ± 3.0% to 9.2% ± 2.9% (corresponding to 82% ± 6.4% reduction) in presence of 5µM 5-NT (Fig 1C). The 50% effective concentration (EC_50_) i.e. the concentration in which 50% reduction is achieved was found to be 2.8µM 5-NT (95% CL, 2.2-3.6 µM). In line with these results, we also observed a reduction in infectious virus particle production. At a concentration of 5µM 5-NT, infectious virus particle production was reduced with >1 log_10_ (94% ± 4.4%) when compared to the vehicle control (Fig 1D). These results correspond to the findings of Mainou and colleagues and confirm that 5-NT interferes with productive CHIKV infection (22).

**Fig 1.**
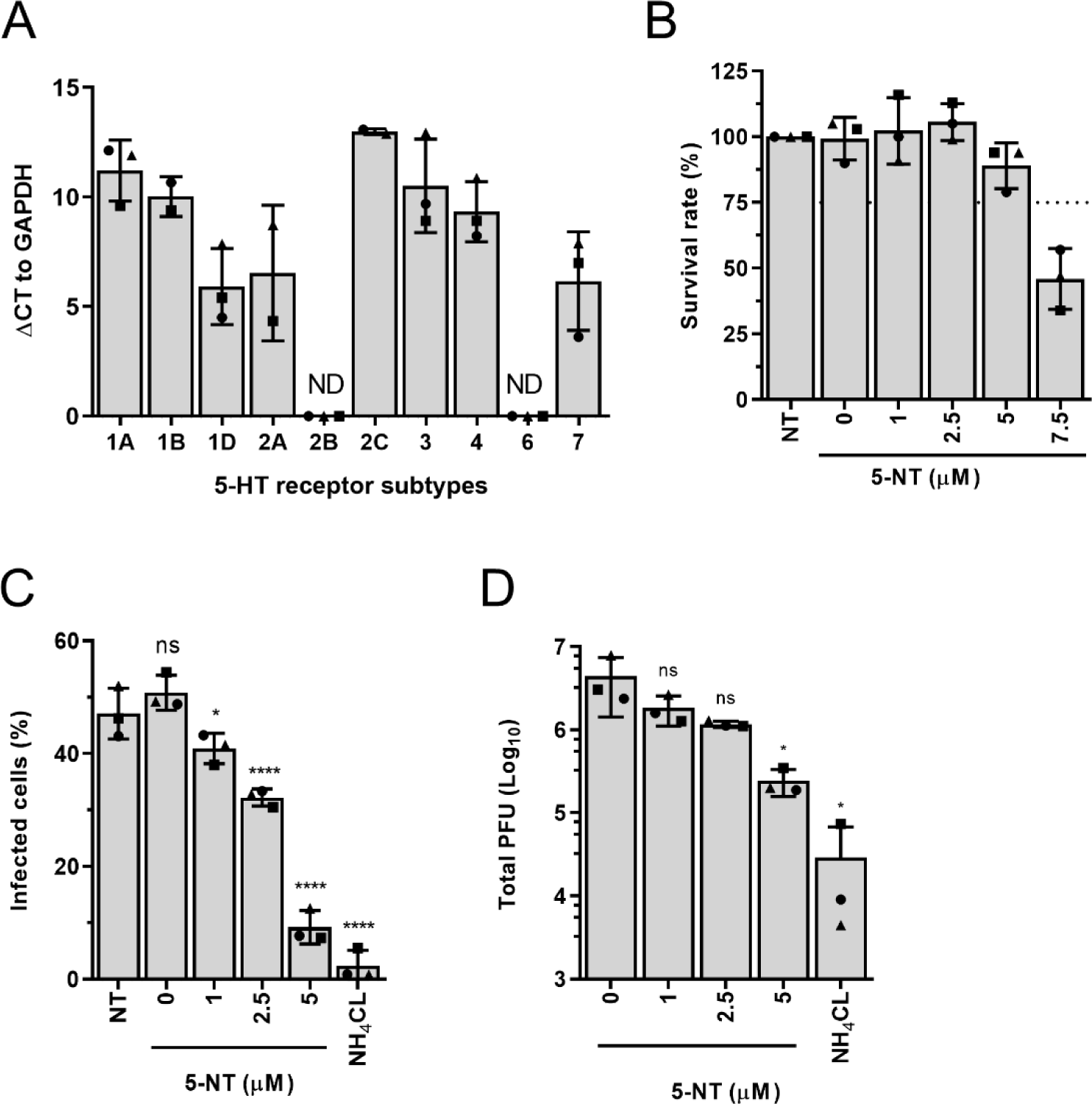
Serotonin receptor agonist 5-NT strongly inhibits CHIKV infection. (A) Delta Ct values between 5-HT receptors and GAPDH mRNA expression in U-2 OS cells. RNA derived from U-2 OS cell lysates was reverse transcribed into cDNA and subjected to qPCR with specific primers for 10 subtypes of the serotonin receptor family and GAPDH (1:10 dilution) as reference gene. Three independent experiments were performed, each in duplicate. Each dot represents the average of an independent experiment. (B) MTT assay to determine the cytotoxicity of 5-NT in U-2 OS cells. Cells were treated for 21 h in the absence or presence of increasing concentrations of the inhibitor to mimic conditions during the infectivity assay. Dotted line indicates 75% cell survival. Three independent experiments were performed, each in sextuplicate. (C, D) U-2 OS cells were pretreated for 1 h with the vehicle control DMSO, 75mM NH_4_Cl or increasing concentrations of 5-NT and subsequently challenged with CHIKV-LR 5’GFP at MOI 5 for 20 h. (B) Cells were collected for analysis with flow cytometry for GFP-positive cells or (C) supernatants were harvested and virus particle production was analyzed by plaque assay on Vero-WHO cells. Three independent experiments were performed. Each dot represents an independent experiment. Bars and error bars represent means and SDs of the experiments, respectively. Statistics was done by use of the student T-test (*P ≤ 0.05). NT, non-treated; ns, non-significant.

### 5-NT inhibits CHIKV infection in first stages of the replication cycle

To unravel the mode of action of 5-NT in controlling CHIKV infection, we first investigated the potential virucidal activity of 5-NT on CHIKV. To this end, CHIKV-LR 5’GFP virions were incubated with 5µM 5-NT for 1.5 h after which viral infectivity was measured by a plaque assay. Within the plaque assay the final end concentration of 5-NT was below 0.5µM as the highest dilution used was 1:10. Incubation with 5-NT did not reduce viral infectivity, demonstrating that 5-NT does not have a direct negative effect on the infectivity of CHIKV particles (Fig 2A). Thereafter, a time-of-addition experiment was performed to delineate where 5-NT acts in the replicative cycle. In these experiments, it is important to analyze the results within one round of replication and therefore we first performed a growth curve analysis on U-2 OS cells. Figure 2B shows that initial GFP fluorescence is detected at 6 hpi (light grey bars, Fig 2B) and robust infectious virus particle production is seen at 8 hpi (dark grey bars, Fig 2B). To increase the sensitivity of the read-out we decided to analyze the effect at 10 hpi which still represents one round of replication. In the time-of-addition assay, cells were treated with 5-NT or DMSO for 1 h prior to infection (pre), during the adsorption of the virus (during), 1.5 h after adsorption (post), or a combination of treatments (Fig 2C). The results were normalized to the DMSO vehicle control. The strongest inhibition of infection was observed when 5-NT was present prior to and during virus adsorption (96% ± 1.2% reduction), which is comparable to treatment during the full course of infection (95% ± 2.6%) (Fig 2D). There is also a clear reduction in viral infectivity when 5-NT was present prior to (71% ± 2.6%) or during (75% ± 1.2%) virus adsorption yet this is significantly lower when compared to complete treatment conditions. Only 29% ± 8.7% reduction in infection is seen when 5-NT is added after adsorption of the virus. Collectively, these data suggest that 5-NT predominantly inhibits CHIKV infection during the early stages of the viral replication cycle.

**Fig 2.**
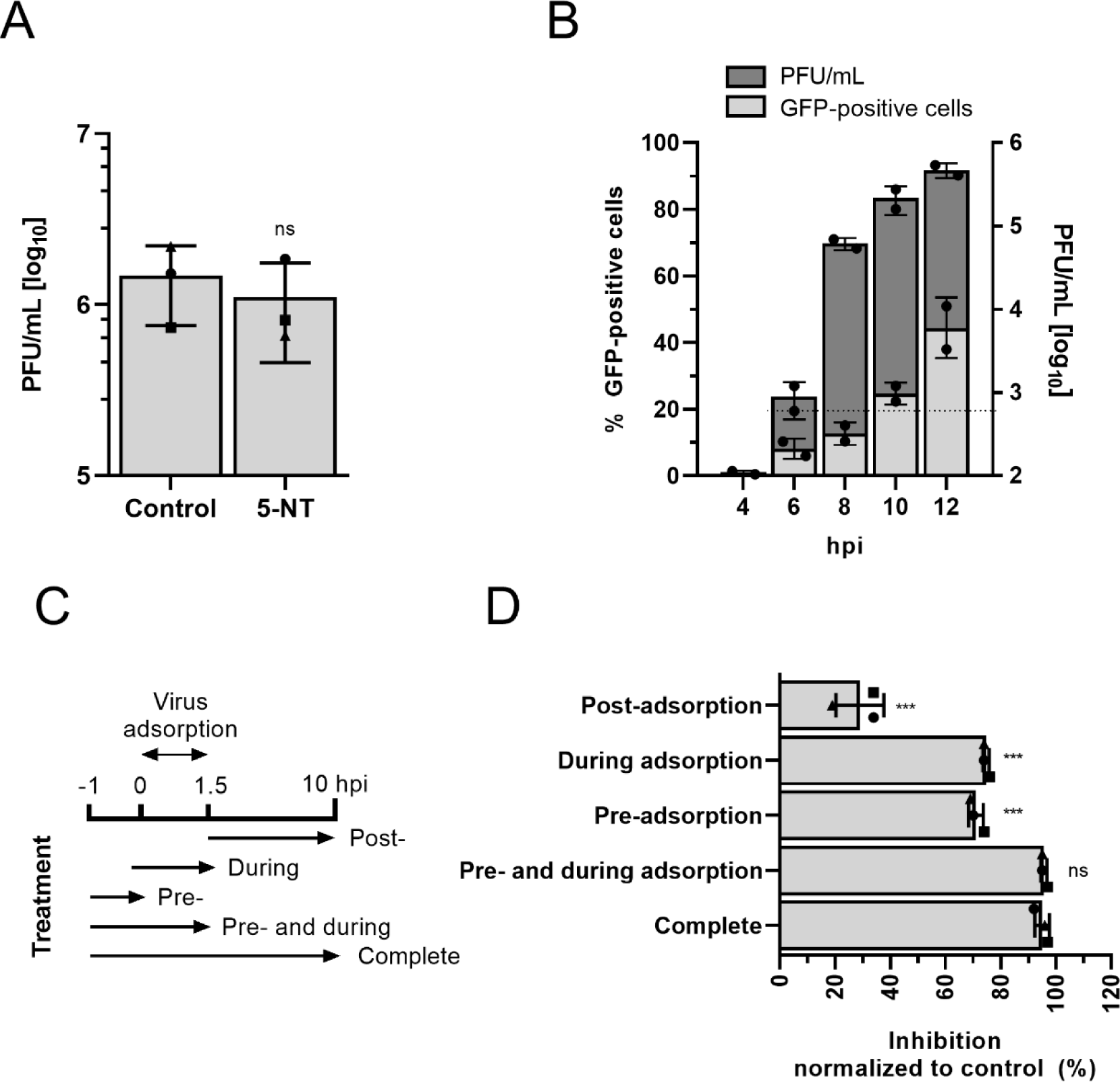
5-NT inhibits CHIKV infection early in the replication cycle. (A) CHIKV-LR 5’GFP was incubated for 1.5 h at 37°C in U-2 OS medium containing 2% FBS and 5µM 5-NT or vehicle control DMSO in a final volume of 300µL. After incubation, the infectious titer was determined by plaque assay in Vero-WHO cells. Three independent experiments were performed. (B) Growth curve analysis of CHIKV infection. U-2 OS cells were infected with CHIKV-LR 5’GFP at MOI 5. Cells and supernatant was collected at 4, 6, 8, 10 and 12hpi to determine GFP-positive cells using flow cytometry (light grey bars) and infectious virus particle production using a plaque assay on Vero-WHO cells (dark grey bars), respectively. Dotted line represents the detection limit of the plaque assay. Two independent experiments were performed, each in duplicate (C) Schematic representation of the time-of-addition assay. (D) U-2 OS cells were treated for the indicated time-points with vehicle control DMSO or 5µM 5-NT. Virus adsorption was allowed for 1.5 h after which the inoculum was removed. U-2 OS cells were collected at 10 hpi and analyzed for GFP-positive cells using flow cytometry. Three independent experiments were performed. The interpretation of each dot, bar, error bar and statistics is explained in the legend to Figure 1.

### Cell entry bypass of the viral genome circumvents 5-NT antiviral activity

To confirm that the serotonin receptor agonist predominantly inhibits CHIKV early in infection, we next evaluated the effect of 5-NT in an infection by-pass experiment. To this end, cells pretreated with 5-NT or vehicle control DMSO were harvested and transfected with *in vitro* transcribed viral RNA by electroporation. Following electroporation, cells were incubated for 12 h in presence of cell culture medium complemented with the compounds. Alternatively, cells were only exposed to 5-NT or DMSO after electroporation. A latter harvesting time-point was chosen to allow for cell recovery due to the electroporation procedure. At these conditions, the infectious virus particle production was 6.6 ± 0.6 Log PFU/mL in DMSO control cells. A comparable virus titer, 6.7 ± 0.2 Log PFU/mL was detected when cells were solely pretreated with 5-NT. Also, no major effect in infectious virus particle production were seen in cells treated with 5-NT at post-electroporation (6.5 ± 0.6 Log PFU/mL) and pre- and post-electroporation (6.3 ± 0.8 Log PFU/mL) conditions (Fig 3A). An inhibitory effect was, however, seen in the percentage of infected cells (Fig 3B). It is important to note, however, that this result might be slightly biased since we detect GFP fluorescence at 6 hpi during normal infection conditions and thus at 12 hpi we may pick-up two rounds of replication. The observed reduction in the number of infected cells may therefore be due to an inhibition in re-infection. Notably, in this experimental set-up, we observed high viral titers, which is indicative for a high transfection efficiency. To validate if 5-NT still exhibits potent antiviral activity at these conditions, we next determined the inhibitory capacity of 5-NT following infection at high MOI. As a control we also determined the viral titer at 4 hpi to ensure that we properly removed the high concentration of virus inoculum and revealed a residual titer of 2.9 Log confirming that we predominantly detect progeny virions at 10 hpi. Under standard infection conditions at MOI 60, a viral titer of 6.0 ± 0.1 Log PFU was observed at 10 hpi (Fig 3C). Importantly, at these conditions, we still observe a robust antiviral activity of 5-NT. The viral titer was 5.1 ± 0.3 Log PFU, which corresponds to 0.9 Log reduction in infectivity in one round of replication (Fig 3C). Altogether, these data confirms that 5-NT predominantly interferes with the early steps of the CHIKV replication cycle, hence before the viral RNA is released in the cell cytosol.

**Fig 3.**
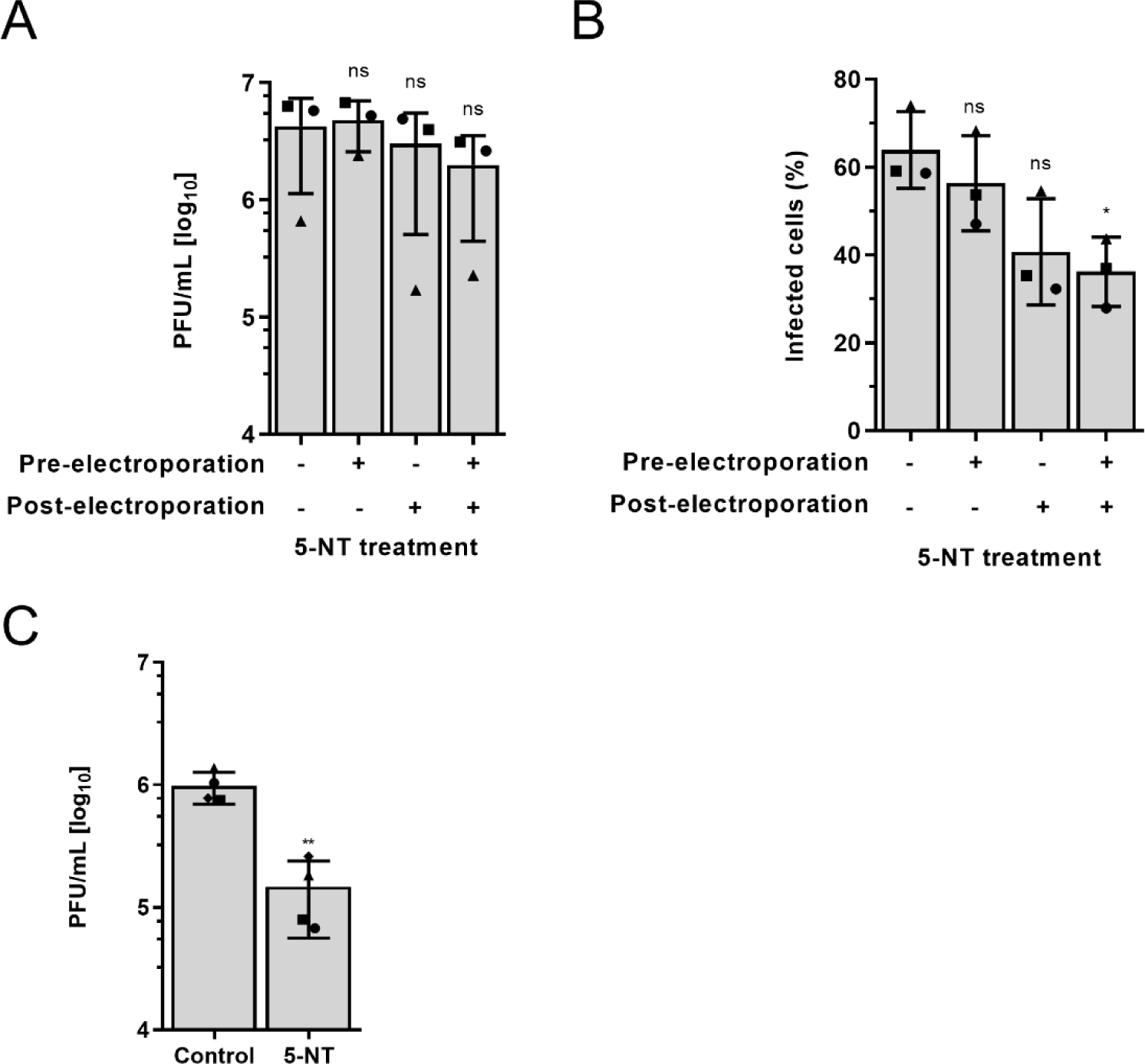
The antiviral activity of 5-NT is before viral genome delivery. (A, B) U-2 OS cells were pretreated with 5-NT or vehicle control DMSO for 1 h before *in vitro* transcribed viral RNA was transfected by electroporation. Cells were cultured for 12 h with or without 5-NT. (A) Supernatants were harvested and virus particle production was determined with a plaque assay on Vero-WHO cells. (B) Cells were collected for analysis with flow cytometry for GFP-positive cells. (C) U-2 OS cells were pretreated with 5µM 5-NT or vehicle control DMSO for 1 h before infection with CHKV-LR 5’GFP at MOI 60. Supernatants were harvested after 10 hpi and virus particle production was determined with a plaque assay on Vero-WHO cells. At least three independent experiments were performed, each in duplicate. The interpretation of each dot, bar, error bar and statistics is explained in the legend to Figure 1.

### 5-NT does not affect CHIKV cell binding

First, we assessed whether the binding capacity and internalization properties of CHIKV in U-2 OS cells is affected in presence of 5-NT. Initially, we determined the interaction of CHIKV with the host cell surface by use of ^35^S-labeled CHIKV particles. To mimic the pre- and during adsorption conditions of the time-of-addition experiments, cells were pretreated with 5-NT or vehicle control DMSO for 1 h after which 1×10^5^ dpm ^35^S-labeled virus (equivalent to ∼1.0×10^9^ GECs and 2.6×10^8^ PFU) was added in cold medium in presence and absence of 5-NT. Incubation was continued for 3 h at 4°C to maximize virus-cell binding. At these conditions, internalization of virus particles is inhibited (31). Thereafter, cells were washed thoroughly to remove unbound virions and harvested by trypsinization. Radioactivity was counted in the total volume by scintillation counting as a measure of virus-cell binding. We measured on average 1.27×10^4^ and 1.33×10^4^ dpm in the absence or presence of 5-NT, respectively (Fig 4A). This indicates that 5-NT does not interfere with virus-cell binding of ^35^S-labeled CHIKV.

**Fig 4.**
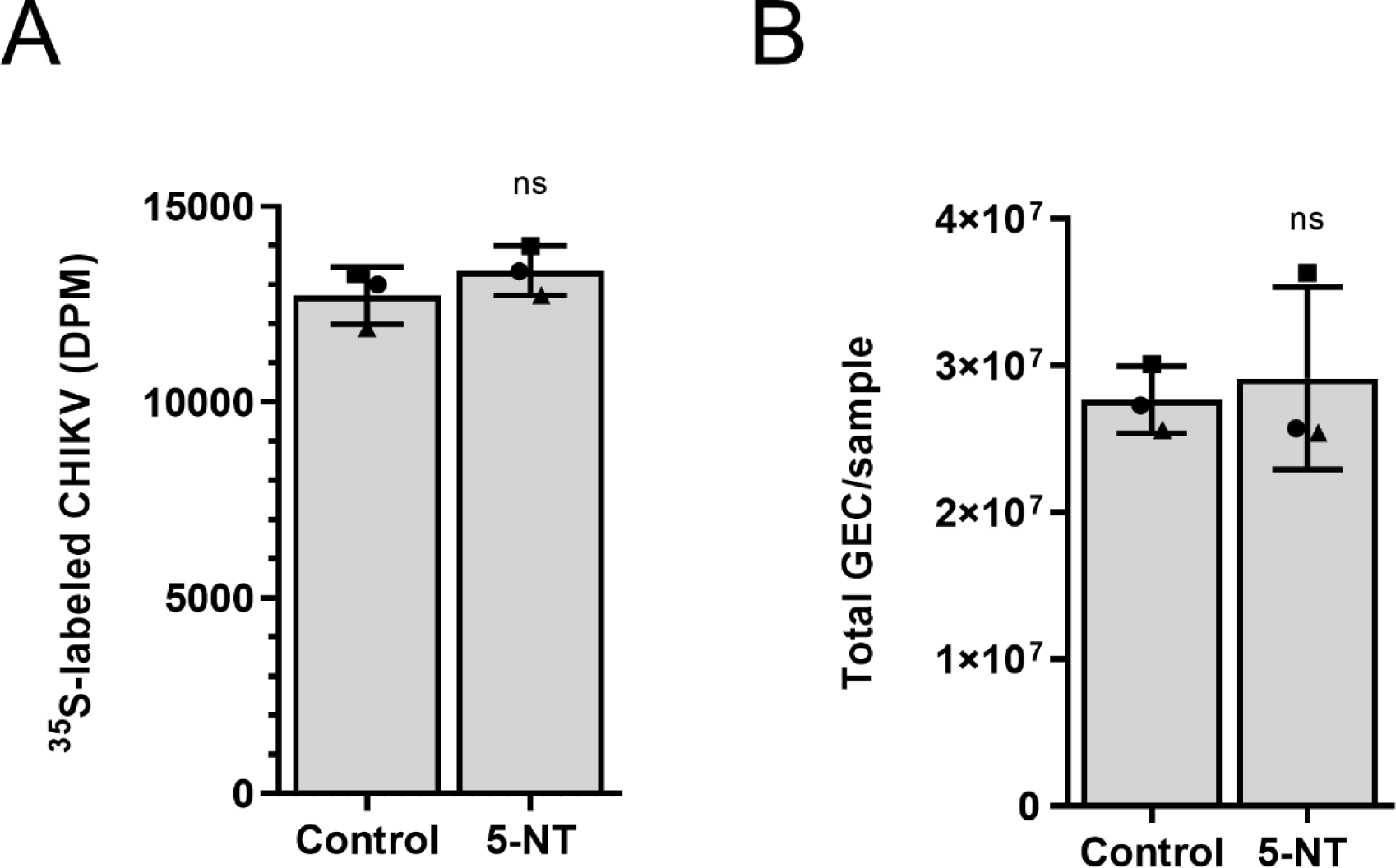
Binding and internalization of CHIKV is not altered in 5-NT treated cells. (A) Binding of CHIKV to U-2 OS cells was tested using ^35^S-labeled CHIKV particles. ^35^S-labeled CHIKV was incubated for 3 h at 4°C in the presence and absence of 5 µM 5-NT with U-2 OS cells. The cells were washed to remove unbound virus particles, harvested and radioactivity was counted. (B) Binding CHIKV particles in 5 µM 5-NT or vehicle control DMSO treated U-2 OS cells. Incubation was done for 1.5 h at 37°C. The cells were washed to remove unbound virus particles and collected to determine total number of viral GECs with specific primers against the viral genome. Three independent experiments were performed, each in duplicate. The interpretation of each dot, bar, error bar and statistics is explained in the legend to Figure 1.

To verify this finding, we next quantified the number of bound and/or internalized GECs by real-time quantitative reverse-transcription PCR (RT-qPCR). To this end, 5-NT or control-treated cells were exposed to CHIKV (∼1.5×10^8^ GECs, corresponding to MOI 5) at 37°C for 1.5 h to allow virus-cell binding and subsequent internalization. We used a shorter incubation time to limit the chance of detecting progeny viral RNA. Furthermore, RT-qPCR is more sensitive compared to the above approach. After 1.5 h, the cells were extensively washed to remove unbound particles and directly lysed in the cell culture plate for RNA isolation and subsequent RT-qPCR analysis. In agreement with the data shown in Fig 4A, we found no difference in total GECs bound and/or internalized between samples treated with 5-NT or vehicle control DMSO (Fig 4B). Collectively, these results indicate that 5-NT does not interfere with CHIKV cell binding at the cell surface.

### Virus internalization and membrane hemifusion is not affected upon 5-NT treatment

Upon internalization, CHIKV traffics through the endosomal pathway towards early endosomes where membrane fusion occurs (12). To assess whether 5-NT interferes with virus internalization and/or membrane hemifusion, we next applied a microscopic virus internalization/hemifusion assay using DiD-labeled CHIKV particles (12, 32, 33). Herein, membrane hemifusion is evident as an increase in fluorescent activity by dequenching of the DiD probe due to dilution within cellular membranes. Membrane hemifusion is a temporary stage prior to fusion pore formation at which the apposed leaflets of the viral membrane and the endosomal membrane have already merged yet the inner leaflets of the lipid membranes are still intact (34). In this assay, the total extent of DiD fluorescence is thus taken as a measure of internalization/hemifusion. U-2 OS cells were pretreated with 5-NT or vehicle control DMSO for 1 h and challenged with DiD-labeled CHIKV particles for 20 min in presence of 5-NT or DMSO. This time-point was chosen as our previous studies revealed that 90% of all hemifusion events occur within the first 20 min post-infection (12). Fig 5A shows representative images for all treatment conditions. As a positive control fusion-inactive DEPC-treated CHIKV was used (12). Quantification of the total DiD fluorescence intensity in 15 randomly selected images revealed that there are no differences in the extent of hemifusion between 5-NT and DMSO treatment conditions (Fig 5B). Taken together, these results suggest that there is no effect of 5-NT on virus cell entry and the membrane hemifusion capacity of CHIKV.

**Fig 5.**
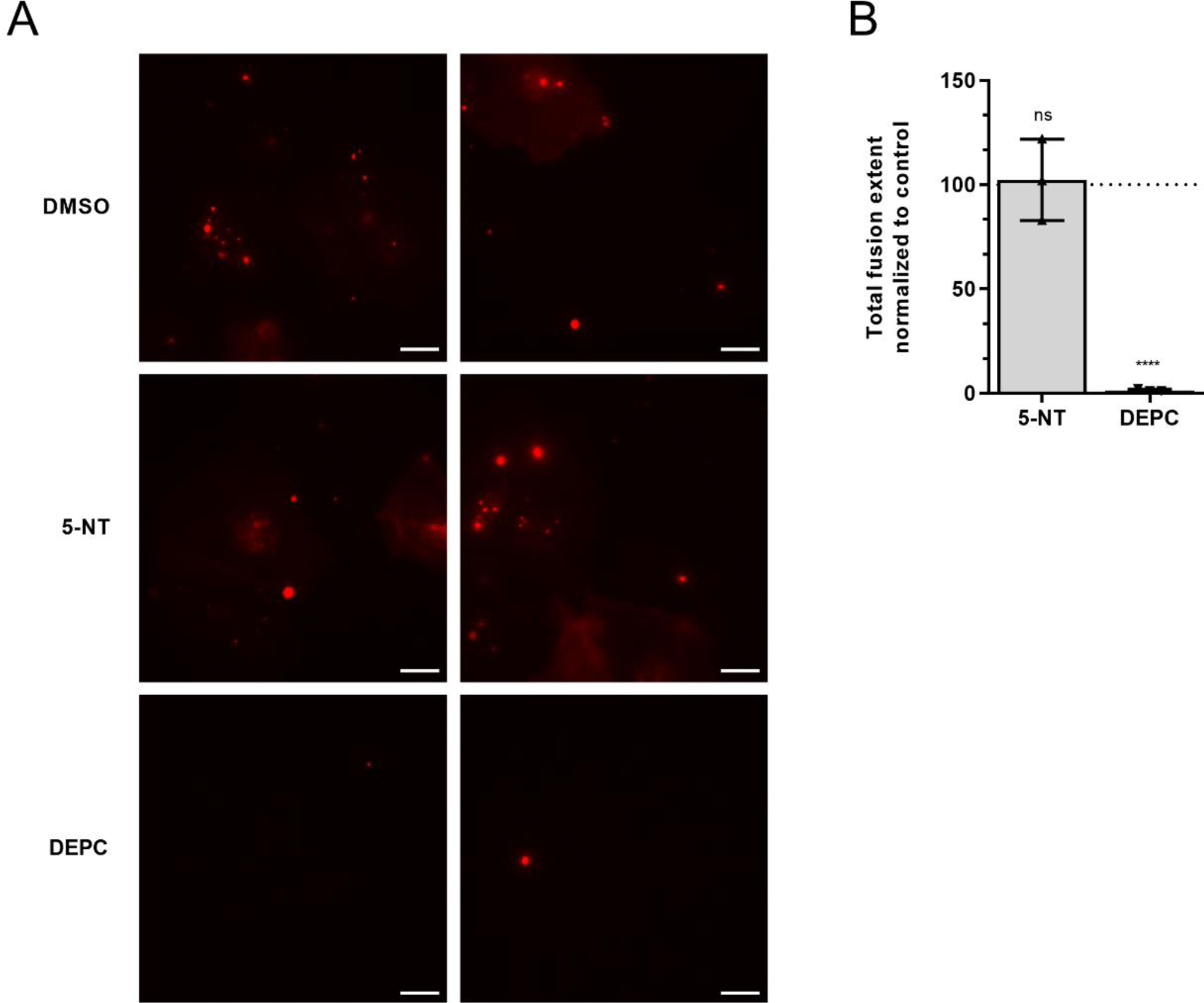
5-NT does not interfere with the membrane fusion capacity of CHIKV. U-2 OS cells were pretreated with 5 µM 5-NT or vehicle control DMSO for 1 h and incubated for 20 minutes with DID-labeled CHIKV particles in presence of 5-NT or DMSO. After washing to remove unbound virus particles, the extent of membrane hemifusion was measured in 15 randomly taken microscopic images per experiment. (A) Representative images showing the DiD signal in DMSO and 5-NT treated U-2 OS cells and for DEPC-inactivated CHIKV. White bar represents 10µm. (B) The total extent of fusion is normalized to that of the non-treated positive control (DMSO). Three independent experiments were performed, each in duplicate. The interpretation of each dot, bar, error bar and statistics is explained in the legend to Figure 1.

### 5-NT treatment does not inhibit fusion pore formation and RNA release from endosomes

To investigate if 5-NT may act at the level of fusion pore formation and nucleocapsid/RNA delivery into the cell cytosol we separated the cytosol from the endosomal membranes by cell fractionation and analyzed the location of the viral genome by RT-qPCR. To this end, U-2 OS cells were pretreated for 1 h with DMSO, 5-NT or bafilomycin A1, an inhibitor of the vacuolar H+ ATPase required for membrane fusion. Subsequently, the cells were incubated with CHIKV at 37°C for 1.5 h in the presence and absence of the compound after which the cells were washed thoroughly. Thereafter, the cells were permeabilized with 50µM digitonin for 5 min at RT and cells were incubated for 30 min on ice to allow cytoplasmic proteins to diffuse into the supernatant. The supernatant (cytoplasmic fraction; indicative for nucleocapsid/RNA delivery) and extracted cells (endosomal membrane fraction; non-fused particles) were collected for RNA isolation and the number of GECs were assessed. In addition, we subjected both cellular fractions to SDS-page and western blotting to verify the efficiency of fractionation (Fig 6A). Herein, we used GAPDH as a marker for the cytoplasmic fraction and the endosomal markers EEA1 and Rab5 were used for the membrane fraction. Fractionation was very efficient with 82% ± 1.4% of GAPDH and 77% ± 9.7% of Rab5 ending up in the cytoplasmic and membrane fraction based on three independent experiments, respectively. Subsequent quantification of the GECs in the membrane and cytoplasmic fraction revealed that bafilomycin A1 treatment abolished RNA delivery into the cytosolic fraction with 0.92 ± 0.04 fold (Fig 6B). Importantly, 5-NT treatment did not interfere with RNA delivery as comparable GECs levels were found in the cytosolic fraction of control-treated cells. In conclusion, 5-NT does not inhibit membrane fusion and the nucleocapsid/RNA is efficiently released from the endosomal membranes into the cytoplasm.

**Fig 6.**
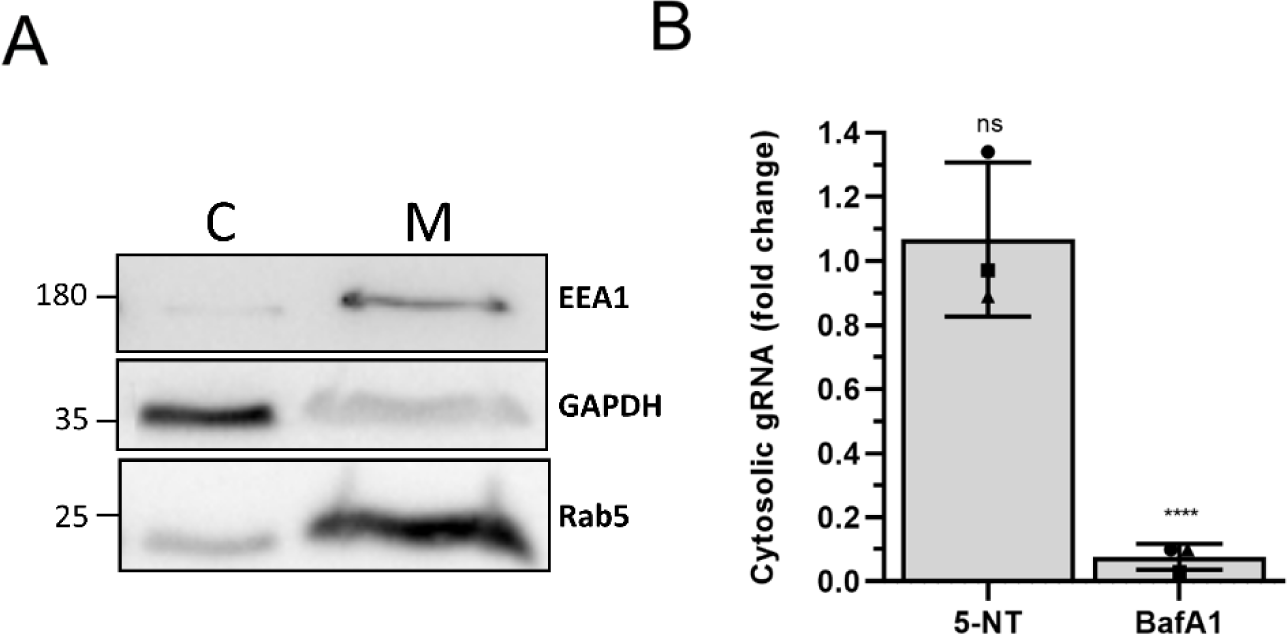
Genomic RNA delivery to cytoplasm upon 5-NT treatment. Pretreated U-2 OS cells with vehicle control DMSO, 5-NT, or Bafilomycin A1 (BafA1) were infected with CHIKV-LR OPY1 2006 at MOI 5 for 1.5 h at 37°C. The cells were washed and permeabilized using 50µg/ml digitonin for 5 min at RT and 30 min on ice. Separation of the cytoplasmic fraction from the membrane fraction was assessed by western blot. Subsequently, the total number of GECs were determined in the cytoplasmic and the membrane fraction. (A) Representative western blot image of the separation. In total three independent experiments were performed. Shown is GAPDH which represents the cytoplasmic fraction (C) and EEA1 and Rab5 which represent the membrane fraction (M) from the extracted cells. (B) Total number of viral GEC was assessed by RT-qPCR after western blot confirmation of successful fractionation. Fold change of cytoplasmic fraction of total gRNA copies is shown. Three independent experiments were performed, each in duplicate. The interpretation of each dot, bar, error bar and statistics is explained in the legend to Figure 1.

### 5-HT receptor antagonist inhibits CHIKV via a different route than 5-NT

The above data shows that 5-NT does not interfere with the initial stages of CHIKV cell entry which is in contrast to what has been described for reovirus (22). In this work the authors also used methiothepin mesylate (MM), which is a 5-HT receptor antagonist blocking 5-HT_1/6/7_ receptors (35, 36) and showed that MM enhanced reovirus infectivity. In an attempt to better understand the above differences we next investigated the role of MM in CHIKV infectivity in U-2 OS cells. First, we assessed the cellular cytotoxicity of MM in U-2 OS cells and revealed that MM is non-toxic to the cells at the concentrations used in this study (Fig 7A). Intriguingly, and in contrast to data described for reovirus, we found a clear dose-dependent reduction in the number of CHIKV-infected cells with 97% ± 1.0% inhibition at 10µM MM (Fig 7B). Due to these contrasting data, we also measured the effect of another 5-HT_1A/1B_ receptor antagonist, isamoltane, on CHIKV infectivity. We chose isamoltane as it has been shown to antagonize signaling pathways downstream of 5-NT at concentrations ranging from 0.01-10μM (37). Our results show that isamoltane is non-toxic to cells at this concentration range (Fig 7C) and does not affect CHIKV infectivity (Fig 7D). This also suggests that the inhibitory effect of MM is independent of 5-HT_1A/1B_ receptor signaling. Importantly, the above data demonstrates that 5-HT agonist and antagonists do not have opposing effects on CHIKV infectivity rather 5-NT as well as MM appear to both act as host-directed antivirals. Given the antiviral role of MM, we next investigated how it interferes with CHIKV infection using similar methods as described above for 5-NT. Time-of-addition experiments revealed that the strongest inhibitory effect (87% ±5.9% reduction) on CHIKV infection is seen when MM is present pre- and during virus adsorption (Fig 7E). Contradictory to 5-NT, pretreatment with MM alone barely inhibited CHIKV infection (20% ± 5.5% reduction), indicating that MM needs to be present during CHIKV adsorption to exert its effect. Indeed, MM treatment during CHIKV adsorption resulted in a reduction of 70% ± 3.7%. Lastly, 30% ± 7.6% reduction was seen when MM was added at post-adsorption conditions. Collectively, the results show that MM, like 5-NT, predominantly interferes with the early steps in infection. Therefore, we next assessed the capacity of CHIKV to bind cells in the presence of MM. In line with the results obtained for 5-NT, no differences in virus-cell binding were observed in the presence or absence of MM (Fig 8A/B). Notably, however, the presence of MM did reduce the total extent of hemifusion activity by 54% ± 11% when compared to the non-treated control suggesting that MM is likely to interfere with internalization and/or membrane hemifusion activity of the virus (Fig 8C). The extent of fusion was comparable to that of cells treated with NH_4_CL (Fig 8C). In line with these results, subsequent cellular fraction experiments revealed that the levels of cytosolic gRNA are reduced to 0.38 ± 0.29 fold compared to the control (Fig 8D). Collectively, this data clearly indicates that 5-NT and MM exert different mechanisms for their antiviral activity in CHIKV replication.

**Fig 7.**
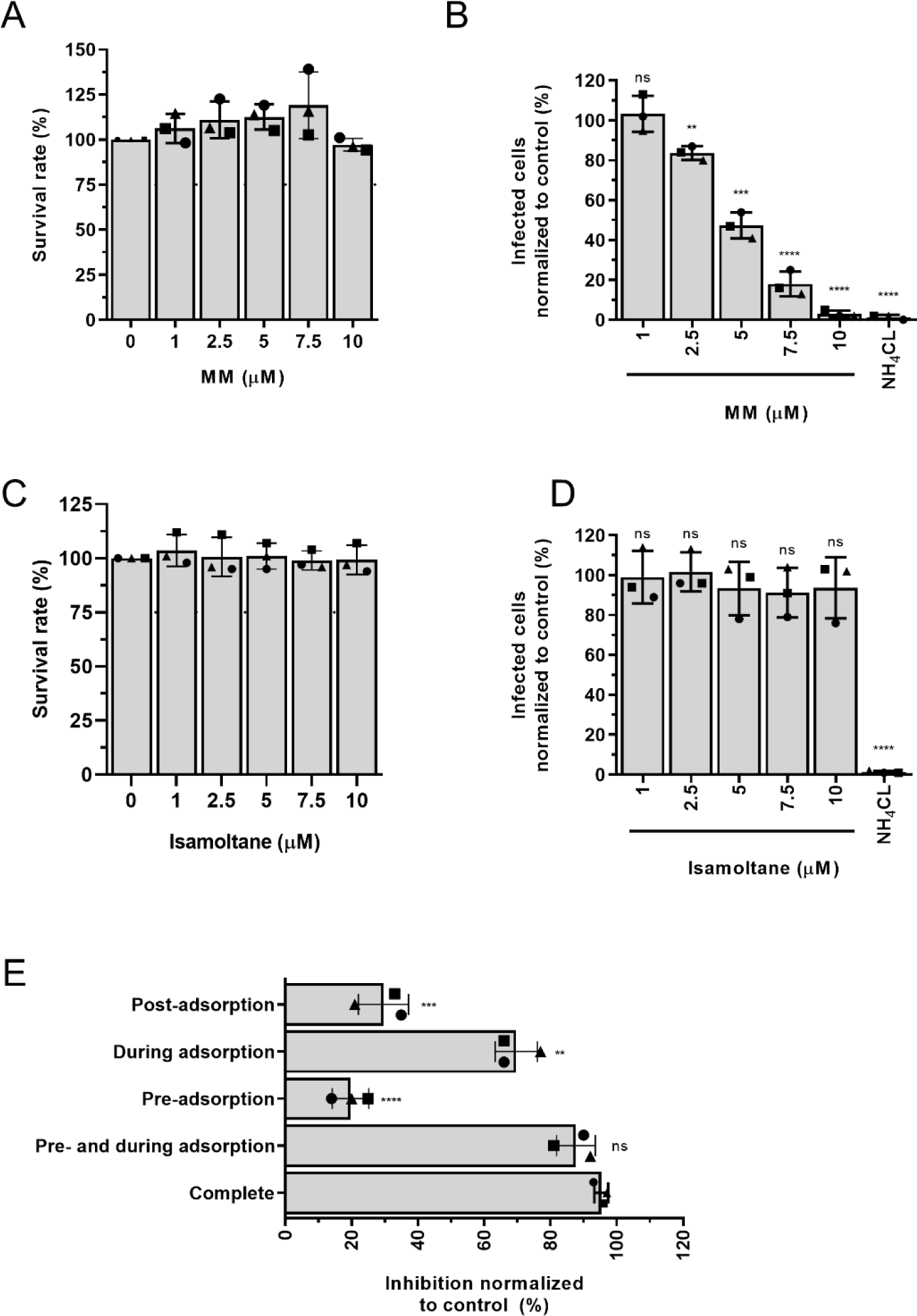
MM inhibits CHIKV infection early in CHIKV infection. (A) MTT assay to determine the cytotoxicity of MM to U-2 OS cells. Cells were treated for 21 h in the presence of increasing concentrations of the inhibitor to mimic conditions during the infectivity assay. Dotted line indicates 75% survival rate. Three independent experiments were performed, each in sextuplicate. (B, D) U-2 OS cells were pretreated for 1 h with 75mM NH_4_Cl or increasing concentrations of (B) MM or (D) isamoltane and subsequently challenged with CHIKV-LR 5’GFP at MOI 5 for 20 h. U-2 OS cells were collected and analyzed with flow cytometry for GFP-positive cells. (C) ATPlite assay to determine the cytotoxicity of isamoltane to U-2 OS cells. Cells were treated for 21 h in the presence of increasing concentrations of the inhibitor to mimic conditions during the infectivity assay. Dotted line indicates 75% survival rate. Three independent experiments were performed, each in duplicate. (E) U-2 OS cells were treated for the time-points indicated in Fig 2A with or without 10µM MM. Virus adsorption was allowed for 1.5 h after which the inoculum was removed. U-2 OS cells were collected at 10 hpi and analyzed with flow cytometry for GFP-positive cells. The interpretation of each dot, bar, error bar and statistics is explained in the legend to Figure 1.

**Fig 8.**
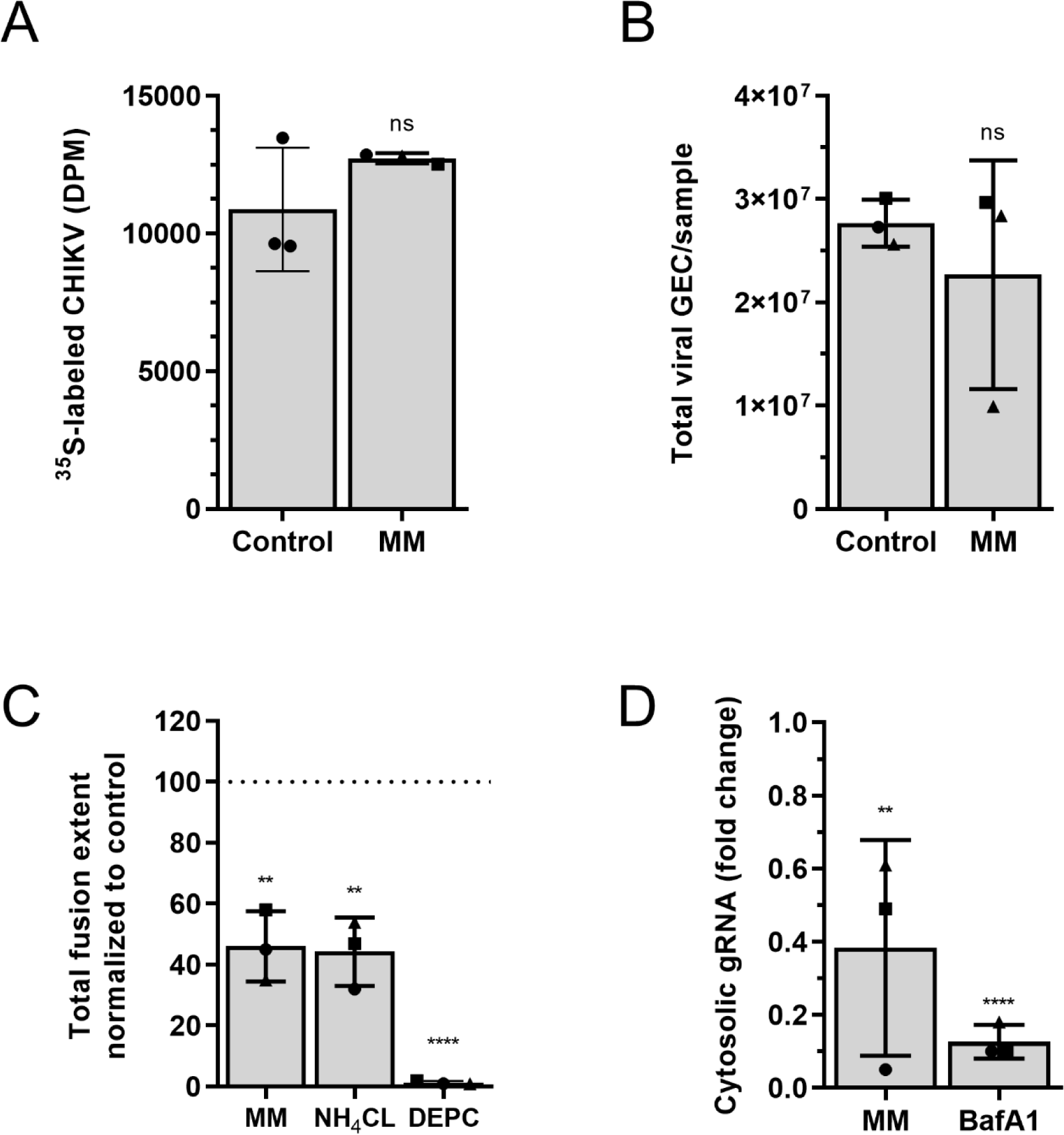
MM inhibits CHIKV infection by restricting membrane fusion activity. (A) Binding of CHIKV to U-2 OS cells was tested using ^35^S-labeled CHIKV. Radioactive labeled virus was incubated for 3 h at 4°C with U-2 OS cells pretreated for 1 h with or without 10µM MM. The cells were washed to remove unbound virus particles and harvested by trypsinization. The trypsinized suspension was subjected to scintillation counting. (B) Binding and internalization of CHIKV particles. Pretreated U-2 OS cells with or without 10µM MM for 1 h were infected with CHIKV-LR for 1.5 h at 37°C. The cells were washed and total number of viral GECs were assessed with specific primers against the viral genome. (C) U-2 OS cells were pretreated with 10µM MM or 50mM NH_4_Cl for 1 h and incubated for 20 min with DID-labeled CHIKV particles in the presence of the compounds. DEPC-inactivated CHIKV was used as a negative control. After washing to remove unbound virus particles, the extent of membrane hemifusion was measured in 15 randomly taken microscopic images per experiment. The total extent of fusion is normalized to that of the non-treated control (DMSO). (D) gRNA delivery to cytoplasm upon MM treatment. Total number of viral GECs was assessed by RT-qPCR. Fold change of cytoplasmic fraction of total gRNA copies is shown. Three independent experiments were performed, each in duplicate. The interpretation of each dot, bar, error bar and statistics is explained in the legend to Figure 1.

## Discussion

In this study we aimed to understand the efficacy and mode of action of serotonergic drugs in CHIKV infection. We focused on the 5-HT receptor agonist 5-NT and 5-HT antagonists, MM and isamoltane. Intriguingly, we observed a strong antiviral effect of both 5-NT and MM on CHIKV infection whereas no effect was seen for isamoltane. We show that 5-NT and MM interfere with distinct steps in the replication cycle of CHIKV.

Addition of 5-NT to cells led to a stark reduction in the number of infected cells and lowered the secretion of progeny virions. Detailed analysis of steps of the replication cycle revealed that 5-NT did not interfere with CHIKV attachment, internalization, hemifusion activity and gRNA delivery to the cell cytosol. Interestingly, however, we also observed that upon transfection of RNA transcripts in 5-NT treated cells, the antiviral activity of 5-NT is almost completely diminished. We have two possible explanations for these intriguing findings. First, even though we do not notice an effect on gRNA delivery to the cell cytosol, we do not know whether the gRNA is still part of the nucleocapsid or not. Thus, based on our findings we hypothesize that 5-NT interferes with nucleocapsid uncoating, thereby reducing the chance to productively infect a cell. The process of nucleocapid uncoating is currently ill-understood and early data suggests that ribosomes are involved in this process (38). However, it has been speculated that as yet unknown host-factors might further contribute to nucleocapsid uncoating (38, 39). Indeed, more recent evidence suggest that ubiquitination and cytoskeleton-associated motor proteins are important for nucleocapsid disassembly in dengue virus, HIV-1 and Influenza A virus infections (40–44). Alternatively, 5-NT stimulation of 5-HT receptors may affect the transport/cellular location of CHIKV-containing endosomes thereby releasing the nucleocapsid at sites that do not support efficient translation and replication of the genome. Indeed, Mainou and colleagues showed that 5-NT treatment did alter the distribution of Rab5 endosomes in CCL2 Hela cells (22).

In this study we also demonstrate that MM behaves as a strong antiviral compound and predominantly controls infectivity after virus cell binding but prior to fusion pore formation and gRNA delivery. Although MM is mainly reported as an antagonist of the 5-HT_1b_ receptor, it also has nonselective properties and can bind to several other receptors subtypes, including 5-HT_6/7_ receptors (35, 45, 46). For example, MM has been described to function as an inverse agonist inducing desensitization of forskolin-stimulated cAMP formation in 5-HT_7_ receptor overexpressed cells (45–47). The lack of antiviral activity of isamoltane strengthens the notion that CHIKV infectivity is not controlled by antagonizing the 5-HT_1B_ receptor. MM and isamoltane are both described as 5-HT_1B_ receptor antagonists yet have distinct alternative effects. For example, isamoltane and MM have been shown to act differentially to the forskolin-induced cAMP formation in renal epithelial cells (48). Future research is required to delineate the precise functions of MM in cells and how it controls CHIKV internalization and/or the process of membrane hemifusion.

Many chemical compound library screen studies have revealed that agonist and antagonist serotonergic drugs can interfere with viral infections (23, 24, 49, 50). For many of these compounds the mechanism of action remains unclear, but many seems to act on cell entry of viruses. For example, in hepatitis C virus infection, 5-HT_2_ receptor antagonists inhibited cell entry at a late endocytic stage. This has been linked to alterations in the protein kinase A (PKA) pathway which interfered with claudin 1, an important receptor for post-binding steps of hepatitis C virus cell entry (17, 49). For JC polyomavirus, 5-HT_2_ receptor antagonists inhibited infection due to interference of binding of β-arrestin to the 5-HT_2A_ receptors, which is required for internalization of the virus via clathrin-coated vesicles (20, 51, 52). During reovirus infection, 5-NT strongly inhibited the cell entry of reovirus whereas MM enhanced reovirus infectivity. This is contradictory to what we observed for CHIKV and this is likely related to differences in the virus cell entry process between both viruses. Reovirus particles traffic towards late endosomes where cathepsin-mediated proteolysis is required for efficient infection whereas CHIKV fusion is solely dependent on low pH and is triggered from within early endosomes (53). Thus, these serotonergic drugs may regulate a host factor that is beneficial for one virus and inhibitory for the other.

The wide spread abundance of serotonin receptors in the periphery and the potent effect of serotonergic drugs on CHIKV infectivity as described in this study suggest that targeting 5-HT receptors might be an interesting approach to alleviate CHIKV disease. Pharmacological targeting of specific 5-HT receptors is, however, challenging due to the various roles of these receptors in multiple parts of the body. To minimize the chance of side-effects it is probably best to use combination treatments with low-concentrations of multiple serotonergic drugs acting on different stages of infection. This will also further reduce the chance of developing resistance to the treatment. Future studies should be performed to investigate the *in vivo* efficacy of single and combination serotonergic drug treatments on CHIKV infection in mice.

## Materials and Methods

### Cells, compounds, and cell viability

Human bone osteosarcoma epithelial U-2 OS cells (a gift from the department of Cell Biology, University Medical Center Groningen, Groningen, The Netherlands) were maintained in Dulbecco’s modified Eagle medium (DMEM) (Gibco, the Netherlands), high glucose, GlutaMAX supplemented with 10% fetal bovine serum (FBS) (Life Science Production, Barnet, United Kingdom). Green monkey kidney Vero-WHO cells (European Collection of Cell Culture #88020401) were cultured in DMEM supplemented with 5% FBS. Baby hamster kidney cells (BHK-21 cells; ATCC CCL-10) were cultured in RPMI medium (Gibco) supplemented with 10% FBS. All media was supplemented with penicillin (100 U/ml), and streptomycin (100 U/ml) (Gibco). All cells were tested Mycoplasma negative and maintained at 37°C under 5% CO2.

Ammonium chloride (NH_4_Cl) (Merck, Darmstadt, Germany) was diluted to a 1M stock concentration in H_2_O. Bafilomycin A1 was diluted to a 200mM stock in dimethyl sulfoxide (DMSO). 5-nonyloxytryptamine oxalate (5-NT) (Tocris, Bristol, United Kingdom) was diluted to a 5mM stock concentration in DMSO (Merck). Methiothepin mesylate (MM) (Tocris) was diluted to a 10mM stock concentration in H_2_O. All chemicals were stored according to the manufacturer’s instructions.

Cytotoxicity of the compounds were tested by use of a MTT [3-(4,5-dimethyl-2-thiazolyl)-2,5-diphenyl-2H-tetrazolium bromide] assay (Merck) at a final MTT concentration of 0.45 mg/ml or by use of a ATPlite Luminescence Assay System (PerkinElmer, Waltham, Massachusetts, United States) according to the manufacturer’s instructions.

### RT-qPCR of serotonin receptors

RNA was isolated from U-2 OS cells with the RNeasy minikit (Qiagen, Hilden, Germany). 0.5µg RNA was reverse transcribed into cDNA using the PrimeScript RT Reagent Kit (Takara, Kusatsu, Japan). Real-time qPCR was conducted on a Stepone plus real-time PCR system from Applied Biosystems using specific primers (Table 1) (Eurogentec, Seraing, Belgium), SYBR green reagents and ROX reference dye (Thermo Scientific, Waltham, Massachusetts, United States). The cDNA was diluted 1:10 for the amplification with GAPDH-specific primers. Data was analyzed using StepOne™V2.3 software.

**Table 1.**
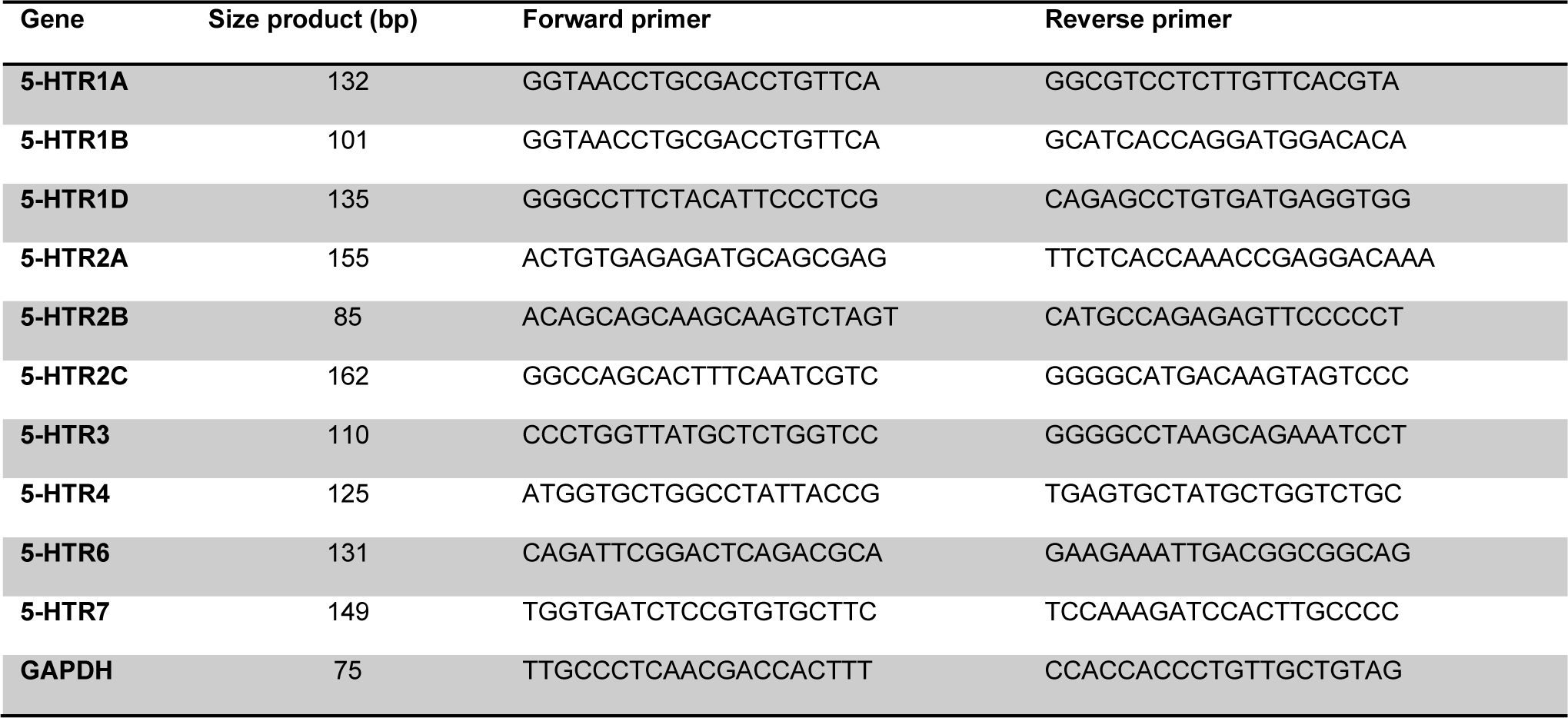
List of primers used for 5-HT receptor genes.

### Virus production, purification and quantification

The infectious clone based on CHIKV strain La Reunion (LR) 2006 OPY1 was kindly provided by prof. Andres Merits (University of Tartu, Tartu, Estonia). CHIKV-LR 5’GFP was kindly provided by the European Virus Archive (EVA, Marseille, France). GFP is cloned after a second subgenomic promotor 5’ to the structural genes (26). Virus production was done as described previously (12, 27). Briefly, BHK-21 cells were transfected with *in vitro-*transcribed RNA transcripts by electroporation with a Gene Pulser Xcell system (1.5 kV, 25µF and 200Ω) (Bio-Rad, Hercules, California, United States). At 22 h post-transfection, the supernatant was harvested (p0) and used to inoculate Vero-WHO cells at a multiplicity of infection (MOI) of 0.01 (p1) to generate working stocks.

Purified virus was prepared by inoculating monolayers of BHK-21 cells with CHIKV-LR (p0) at MOI 4. At 25 hours post-infection (hpi), the supernatant was harvested and cleared from cell debris by low-speed centrifugation. Subsequently, the virus particles were pelleted by ultracentrifugation in a Beckman type 19 rotor (Beckman Coulter, Brea, California, United States) at 54,000xg for 2.5 h at 4°C. The pellet was resuspended overnight in HNE buffer (5mM HEPES (Gibco), 150mM NaCl (Merck), 0.1mM EDTA [pH 7.4] (Merck)) before it was purified by ultracentrifugation on a sucrose density gradient (20 to 50% [w/v] sucrose in HNE) in a Beckman SW41 rotor at 75,000xg for 18 h at 4°C. Upon centrifugation, the virus particles were in the 40% to 45% sucrose layer, which was harvested and aliquoted before storage at -80°C.

L-[^35^S]methionine/L-[^35^S]cysteine-labeled CHIKV was produced by inoculation of a confluent monolayer of BHK-21 cells with CHIKV-LR (p0) at MOI 10. At 2.5 hpi, the BHK-21 cells were starved for 1.5 h with DMEM without cysteine/methionine (Gibco) at 37°C. Next, [^35^S]-EasyTag™ Express Protein Labeling Mix (PerkinElmer) was added and the cells were incubated overnight at 37°C. The medium was harvested and cell debris was removed with low-speed centrifugation. Purification was done by ultracentrifugation for 2 h at 154,000xg at 4°C in a SW41 rotor (Beckman) using a two-step sucrose gradient (20%/50% w/v in HNE). Radioactive virus was collected at the 20%/50% sucrose interface and radioactivity was counted by liquid scintillation analysis. Fractions were pooled based on radioactivity counts and stored at -80°C.

The infectious virus titers of all virus preparations were determined with a plaque assay in Vero-WHO cells. Additionally, the number of genome equivalent copies (GECs) was determined by RT-qPCR, as described previously (11).

### Flow cytometry analysis

Flow cytometry analysis was used to determine the number of infected cells. U-2 OS cells were washed and pre-incubated for 1 h with or without compounds diluted in U-2 OS medium containing 2% FBS. Thereafter, CHIKV-LR 5’GFP was added to the cells at the indicated MOI. At 1.5 hpi, inoculum was removed and fresh U-2 OS medium containing 10% FBS was added in the presence or absence of the compound and incubated for a specified time point at 37°C under 5% CO_2_. Upon collection, cells were washed and fixed with 4% paraformaldehyde (PFA) (Alfa Aesar, Haverhill, Massachusetts, United States) and analyzed by flow cytometry. Flow cytometry was performed with a FacsVerse (BD Biosciences, Franklin Lakes, New Jersey, United States) and analyzed with FlowJo vX.0.7.

### Virucidal assay

CHIKV-LR 5’GFP was incubated for 1.5 h at 37°C in U-2 OS medium containing 2% FBS and 5µM 5-NT or DMSO in a final volume of 300µL. After incubation, the infectious titer was determined by plaque assay in Vero-WHO cells.

### Binding assay with ^35^S-labeled CHIKV

U-2 OS cells were seeded to 80% confluency in a 12-wells plate and washed twice with HNE supplemented with 0.5 mM CaCl_2_ (Merck), 0.5 mM MgCl_2_ (Merck) and 1% FBS (HNE^+^). Cells were incubated with HNE^+^ supplemented with the compounds of interest or vehicle control for 45 min at 37°C and subsequently 15 min at 4°C. Next, 1×10^5^ dpm ^35^S-labeled CHIKV (2.6×10^8^ PFU, corresponding to MOI 500) diluted in HNE^+^ was added to the cells and incubated for 3 h at 4°C to allow virus cell binding. Unbound virus was removed by washing two times with HNE^+^. The cells were harvested by trypsinization and the total volume was subjected to liquid scintillation analysis to count radioactivity.

### Binding and internalization assay by RT-qPCR

U-2 OS cells were seeded to 80% confluency in a 24-wells plate and washed three times with HNE^+^ before incubation with HNE^+^ supplemented with the compounds of interest or vehicle control DMSO for 1 h at 37°C. Next, CHIKV-LR was added to the cells at MOI 5 and incubated at 37°C for 1.5 h. Thereafter, unbound virus was removed by washing three times with PBS (Life Technologies, Carlsbad, California, United States). Next, cells were lysed with the RNAeasy mini kit (Qiagen) according to manufacturer’s instructions and the number of GECs were determined, as described before (11). In addition to this protocol, single-stranded RNA was degraded after cDNA synthesis by RNAse A (Thermo Scientific).

### Microscopic fusion assay

For the microscopic fusion assay, purified CHIKV particles were labeled with the lipophilic fluorescent probe 1,1’-dioctadecyl-3,3,3’,3’-tetramethylindodicarbocyanine, 4-chlorobenzenesulfonate salt (DiD) (Life Technologies), as described before (12). U-2 OS cells were cultured to 80% confluency in Nunc™ 8-well Lab-Tek II Chambered Coverglass slides (Thermo Scientific). Upon infection, the cells were washed three times with serum-free, phenol red-free MEM (Gibco) medium and incubated with phenol red-free MEM supplemented with 1% glucose (Merck) and the compounds of interest. After 1 h treatment, DiD-labeled CHIKV (MOI ∼ 10) was added to the cells and incubated at 37°C for 20 min to allow virus cell entry and membrane fusion. Subsequently, unbound particles were removed by washing three times with serum-free, phenol red-free MEM, after which fresh phenol-red free MEM supplemented with 1% glucose was added. Image fields were randomly selected using differential interference contrast (DIC) and 15 snapshots were taken per experiment in both the DIC and DiD channels with a Leica Biosystems 6000B instrument (Leica Biosystems, Amsterdam, The Netherlands). All snapshots were analyzed for total area of fluorescent spots quantified using the ParticleAnalzer plugin of ImageJ. Total fluorescent area was averaged per experiment and normalized to the total fluorescent area of the vehicle control DMSO.

### Cell entry bypass assay

*In vitro-*transcribed RNA derived from the infectious clone CHIKV-LR was electroporated in 1×10^7^ U-2 OS cells treated beforehand with 5µM 5-NT or DMSO for 1 h, using a Gene Pulser Xcell system (250V, 95µF and 186Ω). After electroporation, the cells were seeded into a 12-wells plate and incubated in medium containing 5-NT at an end concentration of 5µM or the vehicle control DMSO for 12 h at 37°C. Cell supernatants were harvested and analyzed for infectious particle production with a plaque assay on Vero-WHO cells. Additionally, the transfected cells were harvested and prepared for flow cytometry analysis. To this end, cells were fixed with 4% PFA, permeabilized and stained with a rabbit anti-E2-stem antibody (1:1000; obtained from G. Pijlman, Wageningen University, Wageningen, The Netherlands) and Alexa Fluor 647-conjugated chicken anti-rabbit antibody (1:300; life Technologies).

### Digitonin-based cell fractionation

Cell fractionation of U-2 OS cells was performed as described previously (28). Briefly, the cells were seeded to 80% confluency in a 12-wells plate, washed 3 times with HNE^+^, and incubated with HNE^+^ supplemented with the compounds of interest or vehicle control DMSO for 1 h at 37°C. CHIKV-LR was added to the cells at MOI 5 and incubated at 37°C for 1.5 h after which the inoculum was removed. Cells were first washed with PBS, treated for 2 min with an high-salt-high-pH buffer i.e., 1M NaCl in H_2_O [pH 9.5], and then washed for another three times with PBS. Next, cells were permeabilized by incubation with 50μg/mL digitonin dissolved in PBS (Sigma-Aldrich, St. Louis, Missouri, United States) for 5 min at RT and subsequently for 30 min on ice. Afterwards, the supernatant was carefully collected to obtain the cytosolic fraction and the permeabilized cells were collected for the membrane fraction. RNA was isolated using the viral RNA kit and the RNAeasy mini kit for the cytosolic and the membrane fraction, respectively, according to manufacturer’s instructions. The number of GECs were determined, as described before (11). In addition to this protocol, single-stranded RNA was degraded after cDNA synthesis by RNAse A. Additionally, western blot analysis was performed to verify the fractionation step. To this end, the fractions were diluted in 4x SDS sample buffer (Merck) and heated at 95 °C for 5 min prior to fractionation by SDS-PAGE. The antibodies used were mouse-anti-EEA1 (1:5000; BD Biosciences), mouse-anti-GAPDH (1:10,0000; Abcam, Cambridge, United Kingdom), rabbit-anti-Rab5 (1:1000; Abcam). Secondary HRP-conjugated antibodies, anti-mouse or anti-rabbit (Thermo Fisher Scientific) were used as recommended by manufacturer. Quantification was done in ImageQuant TL.

### Statistical Analysis

All data were analyzed in GraphPad Prism software. Data are presented as mean ±SD unless indicated otherwise. Student T test was used to evaluate statistical differences. P value ≤0.05 was considered significant with *p ≤ 0.05, **p ≤ 0.01, ***p ≤ 0.001 and ***p≤ 0.0001. EC50, the concentration at which 5-NT reduces virus particle production by 50% is determined by a dose-response curve that is fitted by lon-linear regression analysis employing a sigmoidal model.

## Acknowledgments

This work was supported by the Graduate School of Medical Sciences of the University of Groningen and by a research grant from De Cock-Hadders Stichting of the University of Groningen (grant to E.M.B). The funders had no role in study design, data collection and interpretation, or the decision to submit the work for publication.

